# Nano-Sieve–Enabled On-Chip Concentration and Multiplexed Detection of Raman-Dye–Labeled Nanocubes

**DOI:** 10.64898/2025.12.17.695009

**Authors:** Tyler Ng, Zhaoxi Yang, Xinye Chen, Kyoung-Jin Yoon, Huiyuan Guo, Eun-Yeong Bok, Katie Lao, Yoon Jung Do, Ruoxue Yan, Ke Du

## Abstract

To address persistent challenges in molecular detection—particularly signal variability, low analyte abundance, and poor spectral reproducibility—a portable bead-stacked nano-sieve platform was engineered to enable robust on-chip Raman signal amplification using silver nanocubes (AgNCs) functionalized with Raman-active thiolated ligands. The microdevice integrates deformable elastomeric channels with magnetic bead stacking to immobilize and concentrate AgNCs within confined nanoscopic junctions, thereby generating densely packed plasmonic hot-spot architectures that markedly enhance Surface-Enhanced Raman Scattering (SERS) efficiency. Time-resolved measurements revealed substantial increases in Raman intensity for both 4-mercaptobenzoic acid (4-MBA) and 4-aminothiophenol (4-ATP), outperforming corresponding off-chip assays and maintaining stable enhancement factors over a four-hour operational window. Furthermore, the platform successfully demonstrated multiplexed sensing capability: mixed-analyte experiments containing 4-MBA, 4-ATP, and cysteamine produced clearly distinguishable spectral fingerprints, even under serial dilution and dynamically pulsed sample delivery. Collectively, these results establish the bead-stacked nano-sieve as a versatile, label-free, and scalable diagnostic architecture capable of achieving high sensitivity and spectral fidelity under continuous-flow conditions. Our technology holds significant promise for real-time, non-invasive molecular surveillance, point-of-care diagnostics, and early disease monitoring in resource-limited or field-deployable settings.

## 1. Introduction

Surface-enhanced Raman scattering (SERS) has emerged as a powerful technique for molecular detection, combining ultrahigh sensitivity with inherent molecular specificity.^1^ In SERS, noble metal nanostructures—particularly those composed of gold or silver—support localized plasmon resonances that intensify the Raman scattering of adsorbed molecules by many orders of magnitude, and this intense field enhancement means that even species with inherently weak Raman signatures can be detected at very low concentrations, approaching or sometimes reaching the single-molecule level.^2–4^ Additionally, SERS provides detailed “fingerprint” spectra: each analyte produces a characteristic pattern of narrow peaks that reflect its molecular structure and bonding. This specificity allows for reliable discrimination among closely related compounds even in complex mixtures. Unlike fluorescence, Raman scatter typically exhibits negligible background and minimal spectral overlap, enabling multiplexing without extensive spectral compensation. The non-destructive nature and photostability of SERS signals further allow for prolonged integration times, enhancing detection reliability.^5–6^

These attributes have made SERS attractive for a wide range of applications, including medical diagnostics^7–9^, environmental monitoring^10,11^, food safety^11,12^, and forensic analysis.^13,14^ However, leveraging SERS in practical settings remains challenging due to the technique’s strong distance dependence: significant signal enhancement occurs only when molecules reside within a few nanometers of the plasmonic surface^15^. This requirement complicates detection in fluidic environments, where analytes must be brought into intimate contact with the nanostructure to achieve meaningful amplification. Conventional SERS substrates—such as planar films, lithographically produced nanostructures, or colloidal nanoparticle aggregates—often suffer from limited control over nanogap formation, leading to variability in signal intensity and poor reproducibility. Aggregation-induced hot spots form unpredictably, and variations in nanoparticle size, shape, and interparticle spacing significantly affect the electromagnetic field distribution. Moreover, many methods rely on static deposition of analytes onto substrates; these approaches are ill-suited for continuous or in situ sensing, particularly in biological or environmental samples where the analyte concentration may be low and transient.^16^

Thiol-functionalized self-assembled monolayers (SAMs)^17^ and film-over-nanoparticle (FON)^18^ strategies have been employed to position analytes close to metallic surfaces, but such techniques can struggle with stability and reproducibility, especially under flow conditions. Additional chemical enhancement mechanisms—such as charge transfer between the metal and analyte—can help boost signal intensities, yet controlling these interactions consistently in a dynamic environment remains a challenge.^19^ The persistent need to pre-concentrate analytes at the surface without compromising reproducibility underscores the importance of developing new platforms that can spatially localize analytes and nanostructures while preserving the sensitivity and specificity of SERS.

To address these issues, we have developed a bead-stacked nano-sieve platform designed for on-chip SERS sensing within a microfluidic environment. This platform integrates deformable microchannels with a packed array of magnetic beads to create a three-dimensional porous medium.^20–25^ By tuning the height and width of the channel to nanoscale dimensions (approximately 200 nm), hydrodynamic forces trap nanoparticles upstream of the bead bed while allowing smaller molecules to pass. Introducing ligand-functionalized silver nanocubes (AgNCs) into this system results in dense, ordered assembly as the cubes become trapped between the beads and within the nano-sieve. The spatial confinement promotes the formation of numerous closely spaced nanogaps, generating abundant “hot spots” for electromagnetic enhancement.^26–27^ The AgNCs can be functionalized with a variety of Raman-active ligands, enabling selective detection of different analytes.^27^ Because the beads are magnetic, they can be easily packed or removed and repositioned, giving the device flexibility and facilitating cleaning or regeneration.^28^ Operationally, the device functions as a continuous-flow concentrator: analytes in the sample stream encounter and interact with trapped AgNCs while being confined within the nano-sieve region. As the AgNCs accumulate, they cluster into well-defined assemblies with consistent interparticle distances, enhancing signal reproducibility.^27–29^ Compared with traditional droplet or static deposition methods, this dynamic approach provides improved control over nanoparticle aggregation and significantly enhances sensitivity by concentrating both the plasmonic nanostructures and the analytes in a small volume. The microfluidic design also allows rapid exchange of reagents and real-time monitoring, features that are critical for diagnostics and environmental applications.

In proof-of-concept experiments, we demonstrated that the nano-sieve platform can efficiently trap AgNCs functionalized with 4-mercaptobenzoic acid (4-MBA), 4-aminothiophenol (4-ATP), and cysteamine. When a dilute suspension of these nanocubes flowed into the device, they accumulated within the bead matrix and formed dense networks, producing strong SERS signals. The detection sensitivity increased markedly compared to off-chip methods, and the signal remained stable for hours, indicating that the microfluidic environment maintained robust hot-spot formation. By introducing different analytes in succession or in mixture, we observed distinct spectral signatures corresponding to each ligand, demonstrating the device’s multiplexing capability. Even under incremental dilution and sequential delivery, characteristic peaks were clearly resolved with minimal cross-interference, illustrating the platform’s potential for analyzing complex samples.

Beyond enabling sensitive detection, the bead-stacked nano-sieve is highly adaptable. Multiple channels, each packed with beads of different sizes or functionalized with distinct ligands, can be integrated onto a single chip for parallel analyses. This modularity lends itself to high-throughput screening and multi-analyte monitoring, which are invaluable in contexts such as point-of-care diagnostics, where rapid identification of multiple biomarkers is essential.^29^ Additionally, by carefully engineering the channel geometry and bead composition, the platform could be tuned for different flow rates, analyte sizes, or environmental conditions, broadening its applicability across various disciplines.

## 2. Materials and Methods

As displayed in **Figure 1a**, fabrication of the nano-sieve channels was developed by bonding a polydimethylsiloxane (PDMS) layer to a 4-inch glass wafer substrate to seal the microchannels for downstream applications. The glass wafer was coated with a ∼200 nm layer of tetraethyl orthosilicate (TEOS) through plasma-enhanced chemical vapor deposition (PECVD). Fabrication for current nano-sieves started with spin-coating a positive photoresist (S-1800 series) onto the TEOS-coated wafer, followed by a dual lithography process. This process served two primary functions: wet etching of the TEOS layer and patterning a ∼1 µm-thick sacrificial photoresist (PR) layer. A UV mask aligner was then used during standard photolithography to define the channel dimensions (12 mm × 2 mm). After UV exposure, the channels were developed with AZ 300 MIF developer, rinsed with water, and then air dried. A 3 mm-thick PDMS layer was then aligned and permanently bonded to the patterned glass substrate through oxygen plasma treatment. After the plasma bond, two holes for the inlet and outlet ports were punctured using a biopsy punch to enable sample loading through microfluidic tubing.

**Figure 1.**
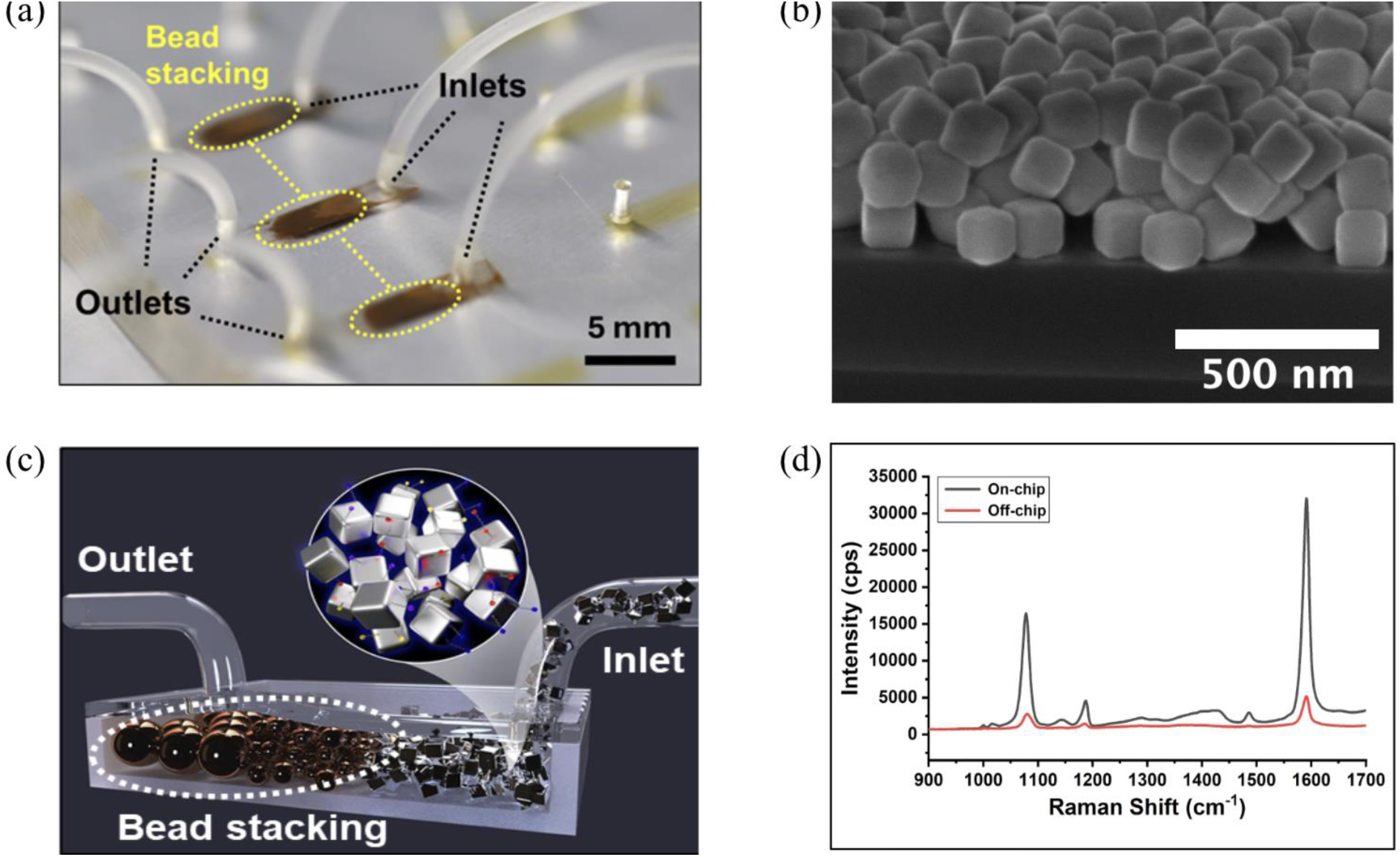
(a) Photograph of nano-sieve channels packed with magnetic beads for AgNCs stacking and concentration. (b) SEM image of Silver Nanocubes used for SERS measurements. (c) Schematic of SERS hot spots formed in a nano-sieve channel. (d) Mean spectra from off-chip (red) and on-chip (black) 4-MBA on AgNC samples, showing significant signal improvement.

Prior to use, the nano-sieve channels were injected with acetone, which completely dissolved the sacrificial PR layer. followed by rinsing the entire channel with isopropanol (IPA) and DI deionized to fully clear the channels. The uncoated magnetic beads with sizes of 10 μm and 1 μm were loaded in the microfluidic tubing connected with a needle and syringe. A syringe pump was then used to inject the magnetic beads into the nano-sieve channel under a flow rate of 15 μL/min for 10 μm beads and 10 μL/min for 1 μm beads, respectively. These beads were essential to the creation of hot spots of silver nanocubes ligand bonded to 4-MBA, 4-ATP, and cysteamine for signal amplification.

In this work, 110 nm silver nanocubes (AgNCs) suspended in ethanol (EtOH) were used as the SERS-active target to validate the nano-sieve sensing platform. As visualized in the SEM image (**Figure 1b**), AgNCs exhibit a natural tendency to aggregate due to both ligand-induced interactions and nanoparticle coupling. However, in off-chip environments, this aggregation is often random and spatially inconsistent, leading to poorly controlled hot spot formation and signal variability.^27^ In contrast, the nano-sieve platform enables on-chip, flow-guided concentration of AgNCs upstream of a bead-packed matrix, promoting localized and reproducible clustering within a confined sensing region. This controlled assembly significantly enhances plasmonic hot spot density and Raman signal consistency. As shown schematically in **Figure 1c**, this effect is further amplified by the use of sulfur-based Raman dyes—specifically 4-mercaptobenzoic acid (4-MBA), 4-aminothiophenol (4-ATP), and cysteamine—which bind strongly to AgNC surfaces via Ag–S bonds, enabling robust ligand exchange and strong Raman signal generation from surface-bound analytes.^25,27^

The nano-sieve is essential for directing AgNC clustering into confined regions, ensuring consistent hot spot formation and reliable SERS enhancement. As shown in **Figure 1d**, the integration of nano-sieve channels resulted in significantly enhanced signal intensities compared to off-chip analysis. This enhancement is attributed to the clustering of AgNCs within the nano-sieve, which facilitates the formation of plasmonic “hot spots” through nanoparticle aggregation. The silver nanocubes (AgNCs) used in this study were synthesized from a modified polyol method adapted from Andrea Tao et al..^30^ In this approach, silver nitrate (AgNO₃) and polyvinylpyrrolidone (PVP) were used as precursors, with 1,5-pentanediol (1,5-PD) serving as both a solvent and reducing agent. To prepare the precursor solutions, 0.20 g of AgNO₃ was dissolved in 10 mL of 1,5-PD alongside 40 µL of CuCl₂ (30 mM), added through sonication in an ice-water bath to ensure uniform dispersion. A separate PVP solution was prepared by dissolving 0.20 g of PVP in 10 mL of 1,5-PD. Both precursor solutions were stored at 4°C until use. For the synthesis, 20 mL of 1,5-PD was preheated in a 100-mL round-bottom flask using an oil bath under continuous stirring at the target reaction temperature. After 10 minutes, 500 µL of the cold AgNO₃/1,5-PD precursor was rapidly injected into the flask, followed by the dropwise addition of 500 µL of PVP solution using a syringe pump at 1-minute intervals. Next, AgNO₃ and PVP were added alternately at a controlled rate to regulate particle growth and achieve uniform nanocube morphology. The reaction mixture was maintained under these conditions for a set duration to allow complete nanocube formation, after which the solution was refluxed briefly to ensure size uniformity. Finally, the flask was removed from the oil bath and immediately cooled in an ice-water bath to halt the reaction.

To functionalize the nanocubes, AgNCs were first synthesized and purified, then mixed with each Raman dye in a 1:1 volumetric ratio (ethanol-based 2M ligand solution to AgNC ethanol dispersion). The mixtures were incubated at ambient temperature for 2 hours to allow sufficient ligand bonding and exchange. This process ensured that the AgNCs were uniformly coated with each respective dye, enabling selective spectral identification and supporting multiplexed SERS sensing within the nano-sieve system.

Following bead stacking and the introduction of AgNCs tagged with Raman dyes into the nano-sieve channels, SERS measurements were performed using a Thermo Scientific™ DXR3 Raman microscope. A 532 nm excitation laser was used for all experiments.^4^ For the off-chip measurements, on-chip nano-sieve measurements, and incremental AgNC accumulation experiments, the system operated at a 0.1 mW laser power with a 1.1 μm spot size, 50 μm slit, and a spectral resolution of 5.5–8.3 cm⁻¹.^31,32^ These measurements were performed using a 50× objective lens, with a 1 second collection exposure time and a 2 seconds sample exposure time per acquisition. These settings were selected to maximize spatial resolution and ensure precise interrogation of the bead-stacked nanogap regions where AgNCs were accumulated.^33,34^

For the multiplexed SERS experiments, a larger sampling area was required to simultaneously probe multiple Raman reporters distributed across the nano-sieve region. To accommodate this, measurements were performed using a 10× objective lens and an increased its laser power to 0.5 mW, while maintaining a 1 second collection time and a 5 seconds sample exposure time.^31,35,36^ The multiplex experiments also utilized a 532 nm excitation source, ensuring consistent spectral alignment with the single-analyte measurements while improving signal throughput across the broader field of view.^3^

## 3. Results and Discussion

Achieving uniform, reproducible SERS substrates that simultaneously allow concentration of analytes and creation of densely packed plasmonic “hot spots” remains a challenge in microfluidic biosensing. The bead-stacked nano-sieve demonstrated in Figure 1 is designed to address these challenges by leveraging microfluidics to concentrate analytes and arrange metallic nanocubes into controlled, three-dimensional assemblies. By flowing solutions of silver nanocubes and thiolated Raman reporters through a microchannel structured with cross-linked polymer beads, the nano-sieve traps the AgNCs, creating numerous sub-nanometer gaps. Such gaps are known to produce intense electromagnetic hot spots that dramatically amplify SERS signals. The platform’s architecture also allows multiple reporters to be functionalized onto the same nanocubes for multiplex detection. To evaluate the effectiveness of this design, three complementary experiments were conducted: assessing the device’s ability to enhance SERS signals by comparing on-chip and off-chip configurations; tracking time-series SERS signals as AgNCs flowed through the sieve to evaluate signal consistency; and testing the platform’s capacity for multi-molecular detection through mixed ligand functionalization.

To rigorously assess the signal enhancement capabilities of our bead-stacked nano-sieve relative to conventional substrates, we conducted on-chip and off-chip SERS experiments using two well-characterized thiolated reporters: 4-MBA and 4-ATP. Equal volumes of functionalized silver nanocubes and reporter solutions were prepared (500 µL of AgNCs mixed with 500 µL of dye) and diluted to 15 µL per measurement. Off-chip samples were generated by depositing this aliquot onto clean glass slides and allowing it to dry, while on-chip samples were obtained by injecting the same volume into the nano-sieve microchannel where the nanocubes are trapped by the bead matrix.

As shown in **Figure 2a**, for 4-MBA, off-chip spectra recorded at twelve random spots exhibited modest peaks at ∼1,084 cm⁻¹ and ∼1,586 cm⁻¹ —vibrations corresponding to the aromatic C–S stretching and ring-breathing modes —with maximum intensities around the low tens of thousands of counts per second (cps).^37^ By contrast, the on-chip spectra displayed markedly stronger signals at the same wavenumbers, with peak heights approaching ∼70,000 cps across all twelve positions (**Figure 2b**). Box-plot analysis of the band near 1,580 cm⁻¹ revealed that on-chip average intensities were roughly two to three times those of the off-chip values, demonstrating that the nano-sieve consistently amplifies the 4-MBA signal while minimizing spot-to-spot variability (**Figure 2e**).

**Figure 2.**
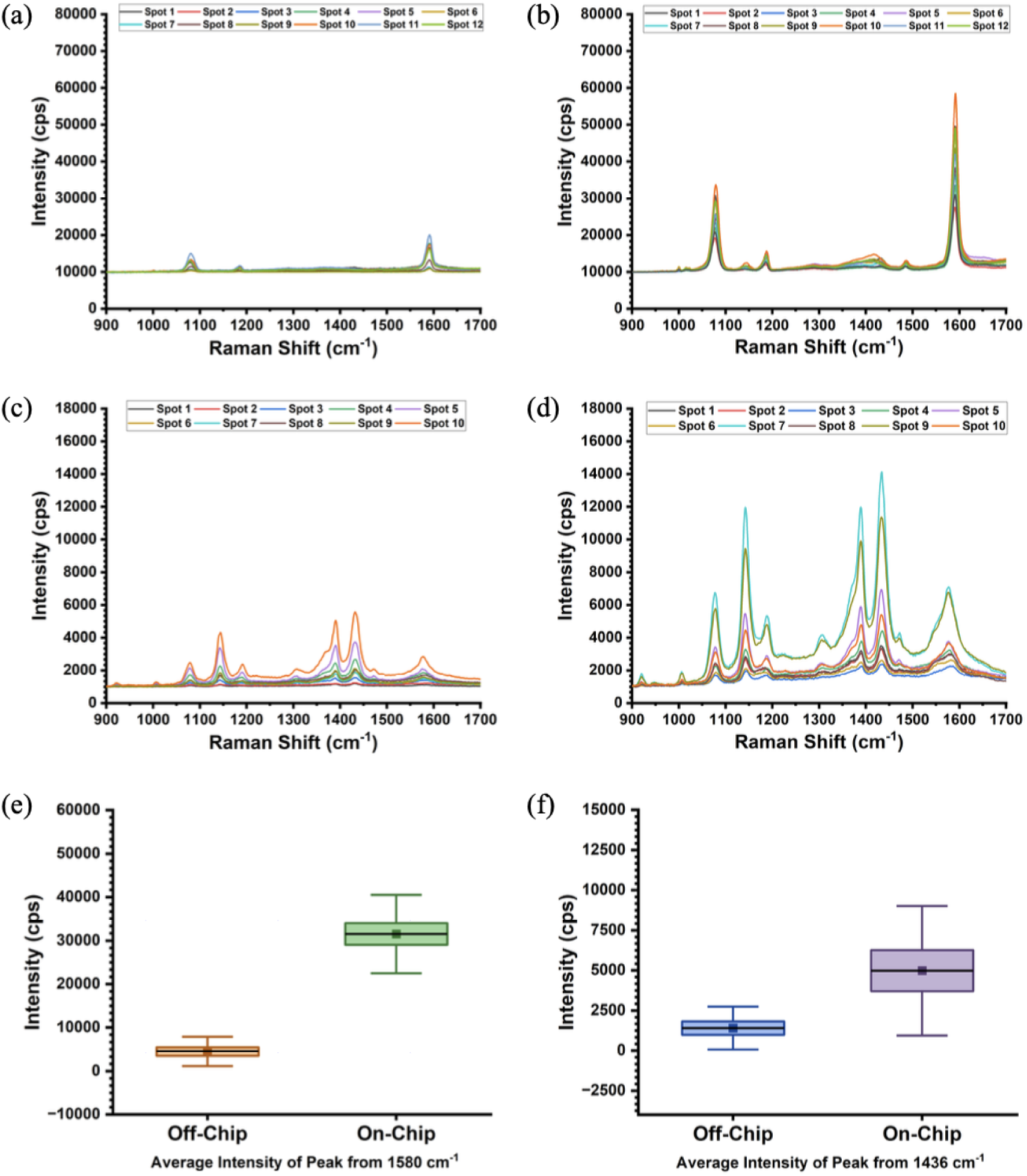
Raman spectroscopy of 4-MBA on AgNC at 12 different random spots: (a) Off-chip and (b) On-chip. Raman spectroscopy of 4-ATP on AgNC at 10 different random spots: (c) Off-chip and On-chip. (e) Mean Intensity of 4-MBA off-chip (Orange) vs on-chip (Green) at Raman shift 1,580 cm^-1^, showcasing significant signal improvement. (f) Mean Intensity of 4-ATP off-chip (Blue) vs on-chip (Purple) at Raman shift 1436 cm^-1^, emphasizing signal enhancement.

A similar trend was observed for 4-ATP. As shown in **Figure 2c**, off-chip spectra from ten spots showed the expected aromatic modes—around 1,084 cm⁻¹ (C–C/C–S), 1,176 cm⁻¹ (C–H), 1,289 cm⁻¹ (C–N), and 1,492 cm⁻¹ —but their intensities were relatively low (typically under 5,000 cps).^38^ When the same nanocube–reporter mixture was introduced into the nano-sieve, all peaks increased sharply, with the 1,436 cm⁻¹ band (used for quantitative comparison) exhibiting an average on-chip intensity more than twice that of the off-chip sample (**Figure 2d and 2f**). These results underscore the nanosieve’s ability to enhance SERS signals across a range of vibrational modes, not just the most prominent ones.

The pronounced improvements observed for both reporters are attributed to the nanosieve’s architecture, which confines AgNCs and analyte molecules within a three-dimensional matrix of polymer beads. This configuration promotes the formation of dense arrays of sub-nanometer gaps—electromagnetic “hot spots”—that dramatically amplify local fields and, consequently, Raman scattering. Furthermore, the consistency in on-chip signal intensities across different sampling locations indicates robust trapping and uniform enhancement, critical for reliable analytical applications. This figure therefore establishes a strong baseline for our device’s SERS amplification performance, setting the stage for subsequent experiments exploring multiplexed detection with additional reporters such as cysteamine.

Following successful validation, subsequent trials were designed to determine whether the device could generate consistent and detectable signals from a diluted nanocube sample delivered at hourly intervals. The nanocube solutions were prepared by ligand bonding 500 µL of silver nanocubes with 500 µL of either 4-MBA or 4-ATP, respectively, followed by dilution in 4 mL of ethanol (EtOH). For both trials, a total volume 150 µL of the diluted nanoparticle solution was extracted and introduced into the nano-sieve device through doses. Specifically, a total injection volume of 37.5 µL per hour was delivered incrementally at a flow rate of 5 µL/min for 7.5 minutes each hour. This resulted in a cumulative delivery of 150 µL into the nano-sieve over a four-hour period.

The following increment trials monitored five locations within the nano-sieve channel for both 4-MBA and 4-ATP samples over a four-hour period. As shown in **Figure 3a**, the 4-MBA spectra collected from a consistent measurement location exhibited a clear time-dependent increase in signal intensity. In particular, the characteristic 4-MBA peaks at 1,078 cm⁻¹, 1,185 cm⁻¹, and 1,590 cm⁻¹ all progressively intensified with each hourly measurement. Quantitatively, the 1,580 cm⁻¹ peak increased from an average of ∼750 cps at hour 1 to ∼6,800 cps at hour 4, corresponding to an approximate nine-fold enhancement. This pronounced rise indicates steady accumulation and clustering of AgNCs upstream of the bead stack, promoting increased formation of SERS-active hot spots within the confined nano-sieve environment. Similarly, **Figure 3b** presents the hourly spectra for a sample labeled with 4-ATP. The prominent 4-ATP peaks at 1,080 cm⁻¹, 1,143 cm⁻¹, 1,390 cm⁻¹, 1,436 cm⁻¹, and 1,575 cm⁻¹ also demonstrated overall upward trends across the four-hour time span. The 1,434 cm⁻¹ peak rose from ∼1,000 cps at hour 1 to ∼1,250 cps by hour 4, representing a ∼25% increase. While the 4-ATP spectra displayed a consistent rise in intensity over time, slight variations, particularly at the one hour mark, may be attributed to minor positional shifts of AgNC aggregates within the nano-sieve during initial flow-in. These shifts can slightly alter local hotspot environments and Raman peak positions. However, the spectral differences remained marginal and did not impede the overall upward trend in signal intensity, reaffirming the platform’s stability. Taken together, the consistent signal increases observed in both dyes confirm the nano-sieve’s ability to continuously trap and concentrate Raman-labeled nanocubes over extended periods, enabling time-resolved signal amplification and enhancing the stability and detectability of SERS signals within the microfluidic channel.

**Figure 3.**
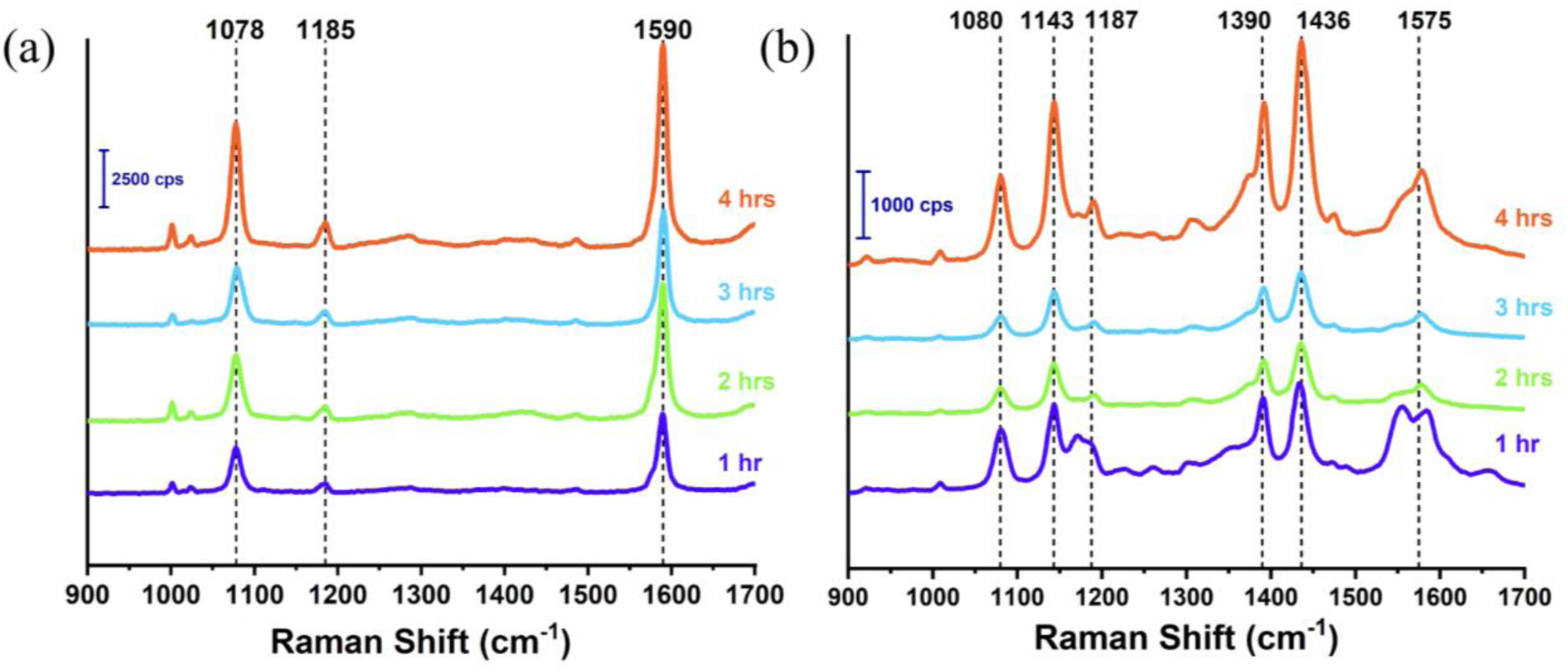
(a) Averaged time-resolved SERS spectra of 4-MBA showing increasing intensity of the characteristic peaks at 1,078, 1,185, and 1,590 cm⁻¹, progressively increase in intensity from the first hour (blue) to the fourth hour (red), indicating continuous accumulation and clustering of AgNCs upstream of the bead stack and the gradual formation of SERS-active hot spots. (b) Averaged time-resolved SERS spectra of 4-ATP, with key peaks at 1,080, 1,143, 1,187, 1,390, 1,436, and 1,575 cm⁻¹, also exhibiting time-dependent enhancement.

To evaluate how effectively the nanosieve could be functionalized with multiple thiolated ligands for multiplex SERS detection, Ag nanocubes (AgNCs) were sequentially modified with three sulfhydryl-binding Raman reporters—4-mercaptobenzoic acid (4-MBA), 4-aminothiophenol (4-ATP), and cysteamine—and the resulting conjugates were mixed before injection into the microfluidic device. Each reporter (100 µL) was allowed to bind covalently to an equivalent volume of AgNCs via thiolate exchange. Two mixtures were prepared to probe the device’s ability to discriminate multiple reporters: (i) an initial mixture containing 15 µL of 4-MBA and 5 µL of 4-ATP to test whether two intense aromatic reporters could be resolved in a mixed SERS spectrum, and (ii) a three-reporter mixture comprising 175 µL of 4-MBA, 25 µL of 4-ATP and 200 µL of cysteamine. The third reporter, cysteamine, is an aliphatic thiol that produces comparatively weak SERS signals; literature shows that its characteristic bands near 630 cm⁻¹ and 712 cm⁻¹ arise from the gauche and trans conformations of the S–C–C chain, so a larger amount was used to ensure detectability.^39^ Each mixture was injected into the nanosieve in 100 µL increments.

SERS spectra collected from three locations along the channel for the first mixture revealed clearly identifiable peaks from both aromatic reporters (**Figure 4a**). 4-MBA produced a strong band at ∼1,084 cm⁻¹ corresponding to an aromatic C-S stretching mode and a prominent ring-breathing band at ∼1,586 cm⁻¹; weaker bands at ∼1,148 and 1,184 cm⁻¹ arise from C-H deformations. 4-ATP (p-aminothiophenol) exhibited intense features at ∼1084 cm⁻¹ (C–C/C–S vibrations), ∼1176 cm⁻¹ (C–H vibration), ∼1289 cm⁻¹ (C–N stretch) and ∼1492 cm⁻¹ (β_CH and C–C modes). Despite the general spectral congruence, the two reporters share vibrational modes around 1,084 cm⁻¹ and 1,180 cm⁻¹, causing their peaks to overlap. Nonetheless, the nanosieve produced reproducible spectra at each sampling spot, demonstrating that surface-bound reporters remained anchored and could be distinguished through their spectral fingerprints.

**Figure 4.**
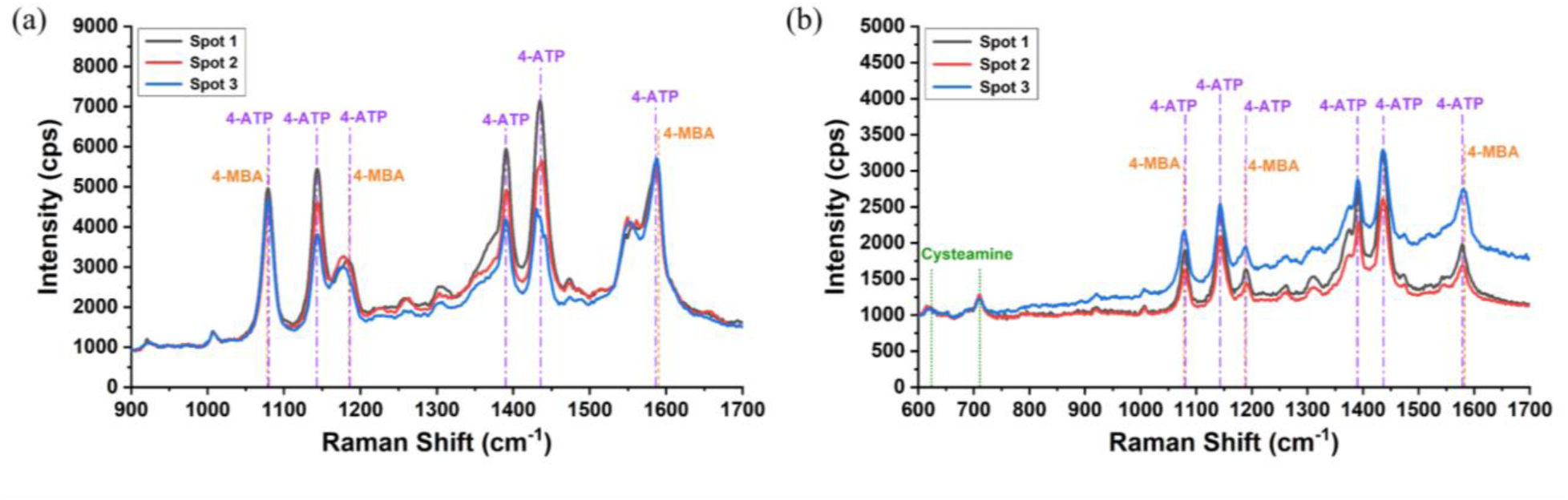
(a) On-chip SERS spectra of a binary mixture containing 15 μL 4-MBA and 5 μL 4-ATP on AgNCs, collected from three spots, showing clearly resolved 4-MBA (1,078, 1,185, 1,590 cm⁻¹) and 4-ATP (1,080, 1,143, 1,187, 1,390, 1,436, 1,585 cm⁻¹) peaks for simultaneous dual-analyte detection. (b) On-chip multiplexed SERS spectra of 175 μL 4-MBA, 25 μL 4-ATP, and 200 μL cysteamine on AgNCs, obtained from three spots, demonstrating concurrent detection of all three analytes through their characteristic Raman signatures, including cysteamine peaks at 633 and 712 cm⁻¹.

In the three-reporter mixture, all characteristic bands were again detected (**Figure 4b**). The cysteamine G- and T-conformer bands at approximately 630 and 712 cm⁻¹ were resolved alongside the intense aromatic features of 4-MBA and 4-ATP. The presence of these aliphatic peaks, despite the dominance of the aromatic bands, underscores the nanosieve’s sensitivity even toward relatively weak Raman scatterers. However, the overlapping vibrational modes of 4-MBA and 4-ATP (most notably the shared C–S stretch near 1,080 cm⁻¹ and C–H deformation region) suggest that peak crowding may limit the number of analytes that can be distinguished using this specific pair of reporters.

Overall, these results demonstrate that the nanosieve microchip can be functionalized with multiple thiol ligands simultaneously, enabling multi-molecular SERS detection within a single device. The successful detection of 4-MBA, 4-ATP and cysteamine across multiple spots confirms that the AgNCs remain surface-bound and retain their spectral integrity. Nevertheless, the overlap between 4-ATP and 4-MBA signals highlights a key design consideration: to fully exploit the multi-ligand capacity of the nanosieve, future studies should explore alternative Raman reporters with non-overlapping spectral signatures. A broader panel of ligands with distinct vibrational modes would improve spectral deconvolution and expand the platform’s multiplexing potential.

Our work demonstrates that integrating a bead-stacked nano-sieve platform with SERS-active AgNCs can significantly improves on-chip Raman signal detection by localizing nanoparticle aggregation and enhancing plasmonic coupling.^26^ Compared to off-chip configurations, which rely on passive aggregation, the nano-sieve enabled a 6-to 7-fold increase in signal intensity for both 4-MBA and 4-ATP, attributed to reproducible hot spot generation within a confined microfluidic space. The controlled environment ensures uniformity in signal enhancement, overcoming one of the critical limitations of conventional SERS substrates, where random clustering often leads to inconsistent signals.

In addition to the 6- to 7-folds signal amplification observed for 4-MBA and 4-ATP, time-resolved experiments further validated the platform’s ability to maintain consistent enhancement over prolonged operation. Incremental SERS measurements taken hourly over a four-hour period revealed a steady increase in Raman intensity, confirming that the nano-sieve can continuously trap and concentrate AgNCs for stable, long-term detection. Furthermore, the successful resolution of distinct spectral peaks from mixtures of 4-MBA, 4-ATP, and cysteamine—despite cysteamine’s relatively weak Raman scattering—demonstrates the system’s ability to detect multiple analytes within a single assay. These findings underscore the reproducibility and sensitivity of the nano-sieve platform, even under dynamic and diluted conditions, and lay the groundwork for its integration into high-throughput microfluidic sensing systems.

The enhanced sensitivity is primarily driven by the physical and chemical confinement achieved through hydrodynamic trapping and magnetic bead stacking within the nano-sieve. This architecture directs AgNC aggregation into densely packed regions, creating stable and concentrated hot spots. Although AgNCs have an inherent tendency to aggregate due to ligand interactions, the nano-sieve allows precise spatial localization and reproducibility, which is not achievable in off-chip conditions.^40^ In addition to the observed signal enhancement, this nano-sieve SERS platform offers a scalable solution for high-throughput sensing due to its compatibility with microfluidic architectures as numerous nano-sieve channels can be patterned on a six inch wafer. The reproducibility and uniformity achieved through spatial confinement distinguish this approach from conventional drop-cast or colloidal aggregation methods, where signal variation can undermine accurate and quantitative analysis.^16^ By introducing programmable flow control, the device could further support kinetic studies of molecular binding events, enabling dynamic monitoring of biochemical interactions in real-time.

While our study establishes a proof-of-concept for nano-sieve-based SERS enhancement, future studies should aim to explore its translation into real-world biosensing applications. Building on the demonstrated multiplexing capacity, the next phase should assess the platform’s performance in detecting clinically relevant biomarkers such as nucleic acids and proteins within complex biological fluids.^21,24,41^ Functionalizing AgNCs with biorecognition elements, such as single-stranded DNA (ssDNA) or antibody fragments will allow for selective and high-affinity analyte targeting.^21,24,41–43^ Moreover, functionalization with disease-specific DNA probes or antibody fragments enables precise recognition of cancer biomarkers, including circulating tumor DNA or protein antigens present in bodily fluids like saliva or urine.^21,44,45^ By enabling sensitive detection of low-abundance molecular signatures in small fluid volumes, this system could be adapted for clinical screening of infectious diseases such as COVID-19, where rapid and reliable identification of viral RNA or proteins is critical.^6,7,44–46^ The platform could be adapted for the detection of protein biomarkers by conjugating AgNCs with aptamers or antibody fragments.^47^ Additionally, optimizing flow dynamics—such as implementing sheath flow—could improve particle focusing and reduce nonspecific accumulation, further enhancing detection sensitivity and repeatability.^48–50^ These developments will not only expand the platform’s diagnostic utility but also contribute to the broader goal of creating robust, miniaturized systems for early disease detection and point-of-care monitoring.

## Acknowledgements

This work is supported by funding from the Rural Development Administration, Republic of Korea for international collaborative Project (PJ017505), USDA National Institute of Food and Agriculture (2024-67022-42826), USDA NIFA (2024-67022-42826), and equipment supplement on National Institute of General Medical SciencesNIGMS (R35GM142763).

